# Correlated variation between the splanchnocranium and basicranium in the Toy rabbit

**DOI:** 10.1101/2022.08.23.504917

**Authors:** Pere M. Parés-Casanova

**Affiliations:** Generalitat de Catalunya, Catalonia, Spain

**Keywords:** brachyfacial morphology, neoteny, palatine morphology, *Oryctolagus cuniculus*, paedomorphy

## Abstract

The aim of this study was to explore and compare patterns of morphological covariation, including symmetrical deviations, between splanchnocranium and the basicranium in Toy rabbit, a type of paedomorphic rabbit. A sample of 32 skulls of adult Toy rabbits was studied on digital pictures on ventral aspects by means of geometric morphometric methods. A set of 7 landmarks were located on the horizontal plane of the splanchnocranial ventral bones (palatine process of the maxillary bone [*processus palatinus maxillae*] + palatine bone [*lamina horizontalis ossis palatini*]), and a set of 8 landmarks were on the basicranium (sphenoid [*os sphenoidale*] + basilar part of the occipital bone [*pars basilaris*] + *bulla tympanica*). Both fluctuating and directional asymmetries were detected on both blocks, being the asymmetry more important among splanchnocranial bones. It appeared also a significant relationship (allometry) between size and shape, especially in the basicranium, as well a significant relationship between the two blocks. From results we deduce that, although normally cranial base usually reaches adult size before the face, in paedomorphic animals the face stops its growth earlier than basicranium, resulting in their proportionally reduced splanchnocranium, the typical brachyfacial morphology for paedomorphy. In other words, our results support the hypothesis of an early stop of facial pattern development in neoteny -the retention of juvenile characteristics into adulthood-, which can affect vital structures. So, further research should expand on clinical data of paedomorphic animals in order to advance in the understanding of pathological results of this growth anomaly.

## Introduction

The bones of the cranial base are known collectively as the basicranium, and include the ethmoid, sphenoid, temporal and occipital bones (Barone, 1999). There we can find the basilar part of the occipital bone [*pars basilaris*] and the sphenoid bone [*os sphenoidale*] (Barone, 1999). This latter is composed by two pieces that can be considered as different bones (Barone, 1999). Basicranium ossifies early during craniofacial development and, being centrally located in the skull, it is thought to act as a structural “foundation” that organizes later growth of the facial skeleton (Adabi, 2018). The ventral part of the splanchnocranium includes, between others, the palatine process of the maxilla [*maxilla*] and the palatine bone [*os palatinum*] (Barone, 1999). They constitute most of the hard palate and floor of the nasal cavity.

Throughout skull development there is a tendency to change skull proportions in two ways: first to increase the size of the brain and second to decrease the size of the jaws, jaw muscles and teeth. But this craniofacial covariation pattern in paedomorphic animals is altered, among whom the overall shape of the skull has been changed: taken together these changes produce a comparatively bigger skull and a shorter face, e.g., a brachycephalia. Toy rabbits present a brachyfacial morphology (Parés-Casanova et al., 2018), adults being much more similar to juveniles than in other rabbit types or breeds, e.g., a paedomorphic appearance, which also includes a flat face, point of ear rounded, a prolonged growth period and a long life span (*pers. obs.*). Toy rabbits’ head can be the same size than the wild rabbit, but different shape. Paedomorphy can be considered a form of neoteny, or the retention of juvenile characteristics into adulthood (Oliver, 2016), a phenomenon observed in many animal species. A paradigmatic example of a neotenous species is the axolotl salamander, which, instead of becoming terrestrial upon reaching sexual maturity, has evolved to retain various juvenile characteristics such as gills and thereby lead an aquatic adult existence.

Among Toy rabbits, the adult facial shape is not very distinct in infancy from adults. Toy rabbits represent an interesting field of study, in part for the understanding of the possible loss of fitness due to this development alteration.

Our aim was to investigate the morphological variability in Toy rabbits on some splanchnocranial and basicranial bones, and specifically, test how asymmetries for both blocks are correlated among this type of animals.

## Materials and methods

### Sample

As the bones are separated by cartilage but ossifies with age, very senile individuals were not sampled due to the difficulty to delimitate clearly sutures. Any of the subjects with a possible presence of craniofacial trauma or malformation were excluded, too. The ultimate sample was comprised of 32 dried skulls of Toy rabbits, without sexual distinction. Ethical and legal requirements were not necessary as they were from natural deaths in the origin farm.

### Imaging

Each skull was photographed on high resolution in standardized ventral view with a Nikon D70 digital camera (image resolution of 2,240 x 1,488 pixels) equipped with a ^®^ Nikon AF Nikkor^®^ 28-200 mm telephoto lens. Scale was given for each photograph by placing a ruler. The images were stored in JPG format.

### Landmark selection

A total of 15 points were chosen to analyse forms. A set of 7 landmarks were located on the horizontal plane of the splanchnocranial bones (palatine process of the maxillary bone [*processus palatinus maxillae*] + palatine bone [*lamina horizontalis ossis palatini*]), and a set of 8 landmarks on the basicranium (sphenoid [*os sphenoidale*] + basilar part of the occipital bone [*pars basilaris*] + *bulla tympanica*) (Figure 1 and Table 1). The *x* and *y* coordinates of these landmarks were digitized using TpsDig 2.04 v. 1.40 (Rohlf, 2015) on images. Digitalizations were made twice in order to assess error. Landmarks were superimposed using Generalized Least Squares Procrustes analysis, which removes information about location and orientation from the raw coordinates and scales each specimen to unit centroid size (CS). CS is defined as the square root of the sum of squared distances of a set of landmarks from their centroid (Webster and Sheets, 2010). The resulting Procrustes shape coordinates and CS were used for statistical analyses.

**Figure 1.**
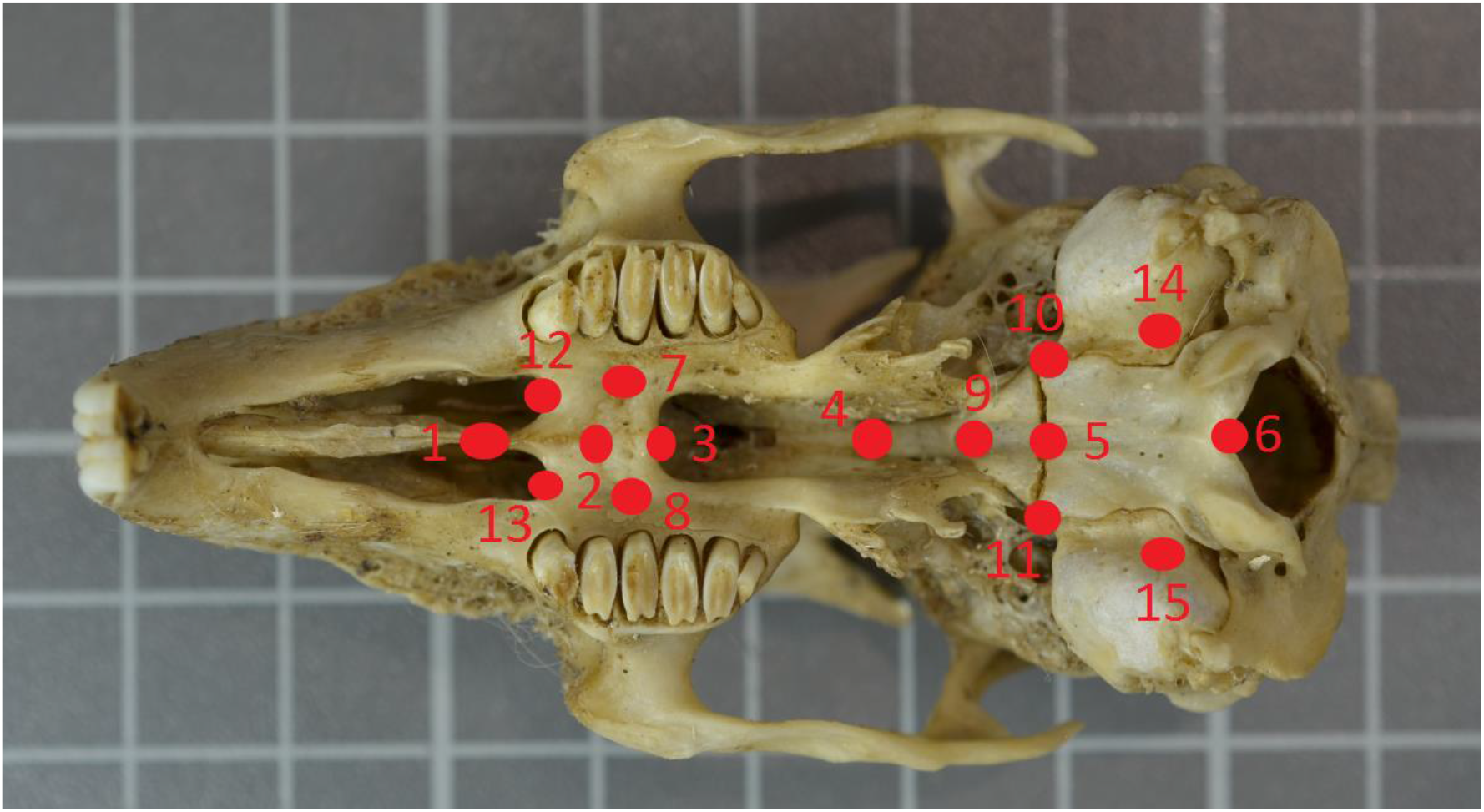
Landmarks locations used it this study on the skull ventral aspect. A set of 7 landmarks were located on the horizontal plane of the splanchnocranial bones (maxillary bone [*processus palatinus maxillae*] + palatine bone [*lamina horizontalis ossis palatini*]), and a set of 8 landmarks on the basicranium (sphenoides + basioccipital+ *bulla tympanica*). See table 1 for a detailed explanation.

**Table 1.**
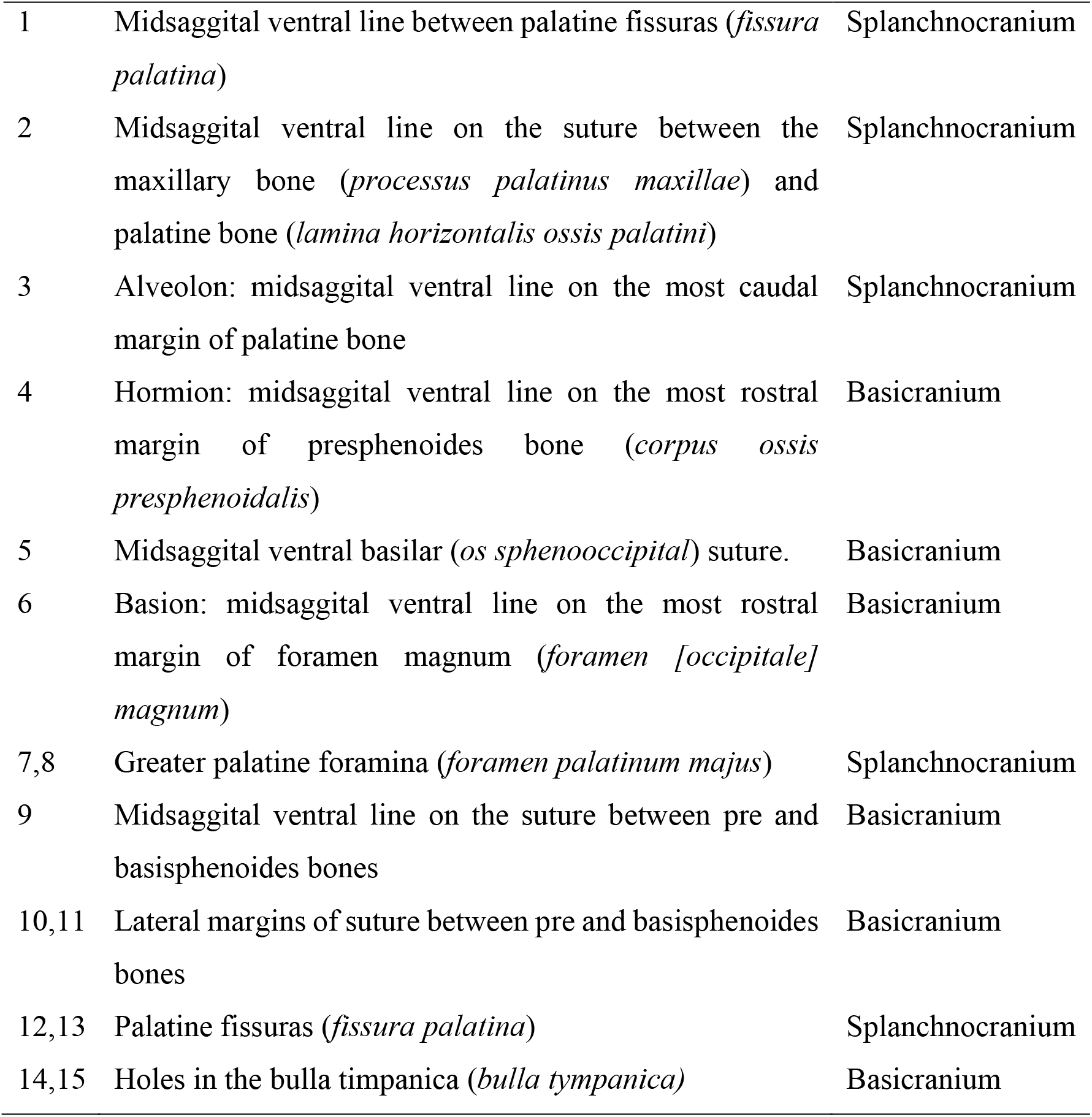
Landmarks used it this study. A set of 7 landmarks were located on the horizontal plane of the splanchnocranial bones (maxillary bone [*processus palatinus maxillae*] + palatine bone [*lamina horizontalis ossis palatini*]), and a set of 8 landmarks on the basicranium (sphenoides + basioccipital).

### Allometry

We used a multivariate regression of shape coordinates for symmetric component on the logarithm of CS for each block for potential influence by allometry, e.g., how differences in size can affect shape. Multivariate regressions were performed independently for the first set of landmarks (splanchnocranium) and for the second set (basicranium) separately.

### Principal Component Analysis

To inspect the variability of landmark points we ran a Principal Component Analysis (PCA) on asymmetric scores performed on all the landmarks for two blocks separately. PCAs were done for both blocks on regression scores, allowing to remove the effect of size.

### Craniofacial integration

Some studies on mammals highlight the covariation of facial size with the rest of the cranium, particularly the cranial base, contributing to the integration and modularity of the skull (Cardini and Polly, 2013). Intraspecific covariance between shapes was assessed by performing Partial least squares (PLS) analyses. The aim of the PLS is to maximize the covariance patterns between two blocks of variables rather than the intra-block variance. PLS describes data in terms of a score for each specimen along a single axis, similar to a principal component that is generated in a PCA. The primary difference is that, unlike principal components, which produces principal axes, PLS produces pairs of axes. Since allometry can inflate measures of integration, the PLS analyses were recomputed using the residuals of the multivariate regression of shape variables on the logarithm of centroid size as variables. We used the RV coefficient to measure the correlation resulting from the PLS. The use of the RV coefficient for the measure of the association between two blocks of variables is recommended in several recent papers, notably because it is calculated directly on covariance and variance rather than on correlation values. The calculation of this coefficient is equivalent to the calculation of the correlation coefficient of a regression between two variables. The RV coefficient is a measure of the global integration between blocks, providing a global quantification of the strength of the association between blocks.

All analysis were done using the MorphoJ v. 1.06c package (Klingenberg, 2011).

## Results

### Measurement error and asymmetries

In the Procrustes ANOVA, where three factors (individual, side and interaction individual*side) were analysed, error was very low (Table 2). Although the interaction of individuals and sides showed a highly significant difference (*p*<0.0001) only for splanchnocranium, Pillai’s trace showed its presence also for basicranium (Pillai’s traces=0.36; *p*=0.0492). Levels of these detected asymmetries (fluctuating asymmetry+directional asymmetry) were much higher in splanchnocranium (69.8%) than in basicranium (19.3%).

**Table 2.**
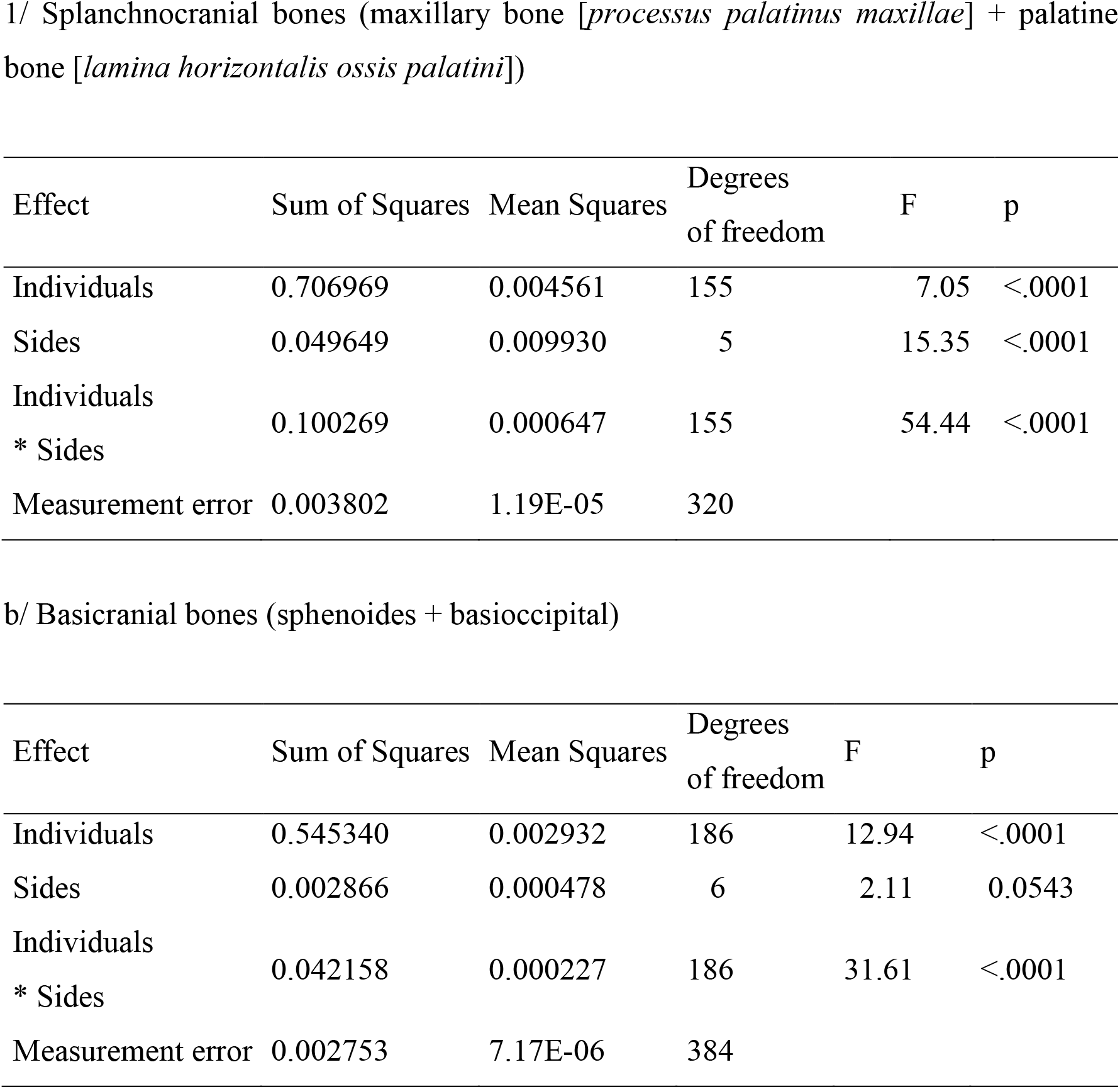
Procrustes ANOVA results for palate and basicranial bones among Toy rabbits (n=32).

| Effect | Sum of Squares | Mean Squares | Degrees of freedom | F | p |
| --- | --- | --- | --- | --- | --- |
| Individuals | 0.706969 | 0.004561 | 155 | 7.05 | <.0001 |
| Sides | 0.049649 | 0.009930 | 5 | 15.35 | <.0001 |
| Individuals<br>* Sides | 0.100269 | 0.000647 | 155 | 54.44 | <.0001 |
| Measurement error | 0.003802 | 1.19E-05 | 320 |  |  |

b/ Basicranial bones (sphenoides + basioccipital)
| Effect | Sum of Squares | Mean Squares | Degrees of freedom | F | p |
| --- | --- | --- | --- | --- | --- |
| Individuals | 0.545340 | 0.002932 | 186 | 12.94 | <.0001 |
| Sides | 0.002866 | 0.000478 | 6 | 2.11 | 0.0543 |
| Individuals<br>* Sides | 0.042158 | 0.000227 | 186 | 31.61 | <.0001 |
| Measurement error | 0.002753 | 7.17E-06 | 384 |  |  |

### Allometry and PCA

The multivariate regressions of Procrustes coordinates of symmetric component (dependant variables) on size (log CS as independent variable) showed a significant influence of allometry for both blocks, being the covariation much higher in basicranium (29.07%, *p*<0.001) than in splanchnocranium (8.75%, *p*=0.001). Data for PCA for splanchnocranial bones is shown in Table 3. According to the results in PC1 and PC2 in splanchnocranial bones, with a total of 76.1% (Table 3), changes were found greatest on landmarks corresponding to most rostral and caudal parts, and palatine foramina (landmarks 1, 3, 7 and 8). For basicranial bones, results in PC1 and PC2, with a total of 81.4% (Table 3), main changes was found mainly on the rostral part of the presphenoides (*os praesphenoidale*), lateral sutures between presphenoides and basisphenoides (body of caudal sphenoid, [*os basisphenoidale*]) and tympanic holes (landmarks 4, 10, 11, 14 and 15).

**Table 3.**
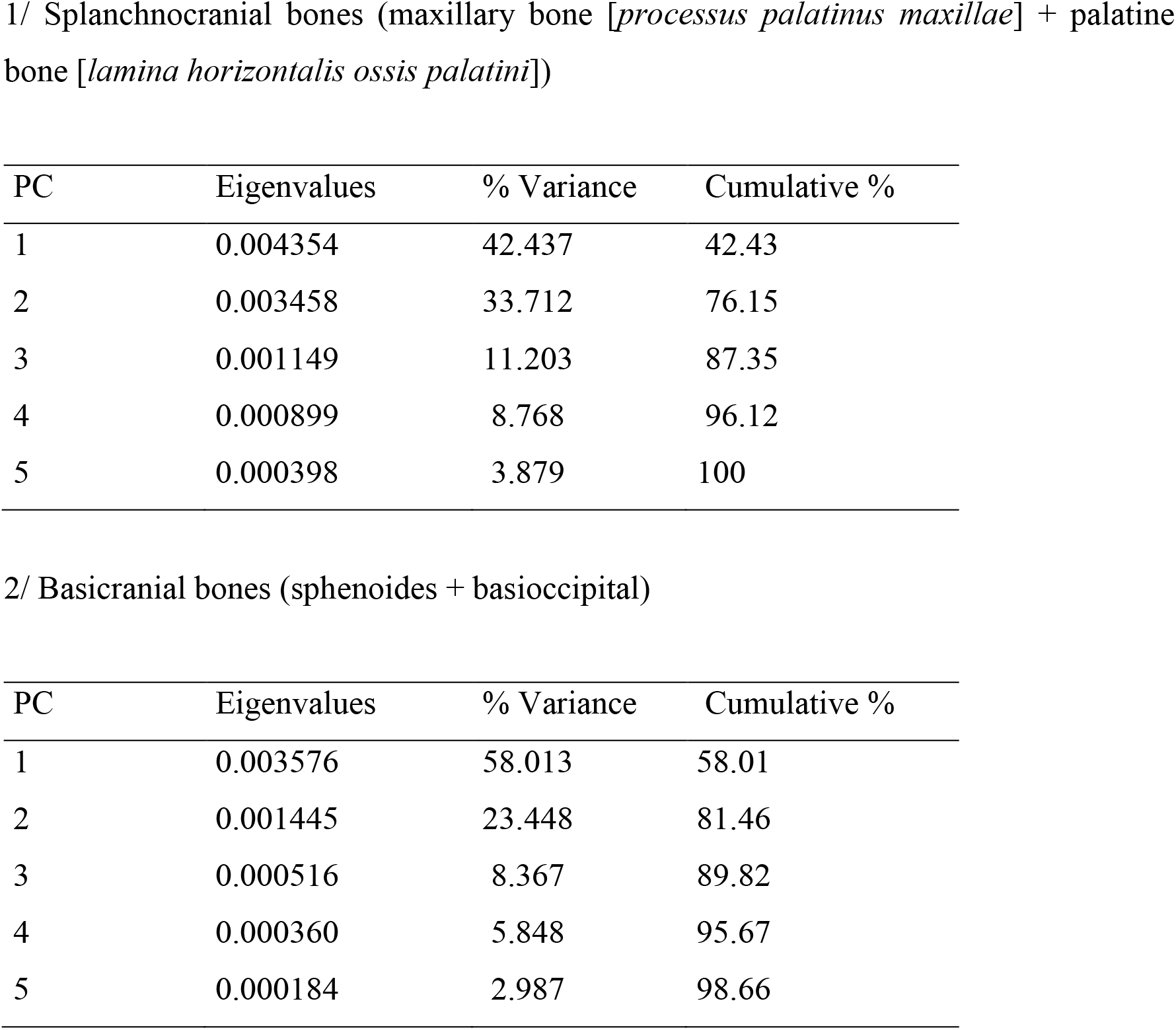
Principal Component Scores for first 5 Principal Components (PC) showing the values for shape scores.

| PC | Eigenvalues | % Variance | Cumulative % |
| --- | --- | --- | --- |
| 1 | 0.004354 | 42.437 | 42.43 |
| 2 | 0.003458 | 33.712 | 76.15 |
| 3 | 0.001149 | 11.203 | 87.35 |
| 4 | 0.000899 | 8.768 | 96.12 |
| 5 | 0.000398 | 3.879 | 100 |

2/ Basicranial bones (sphenoides + basioccipital)
| PC | Eigenvalues | % Variance | Cumulative % |
| --- | --- | --- | --- |
| 1 | 0.003576 | 58.013 | 58.01 |
| 2 | 0.001445 | 23.448 | 81.46 |
| 3 | 0.000516 | 8.367 | 89.82 |
| 4 | 0.000360 | 5.848 | 95.67 |
| 5 | 0.000184 | 2.987 | 98.66 |

### PLS

In the PLS analysis, when the effect of size was removed, the RV coefficient indicated a low although significant relationship between the two blocks (RV=0.174; *p*=0.0168) with the first pair of singular axes presenting 62.5% of the covariance.

## Discussion

The skull is a complex organisation of bones, with morphology being influenced by all its parts. In this study, we aim to assess the morphological relationship between some ventral splanchnocranial and basicranial bones. To achieve this, we used geometric morphometrics on 32 crania of Toy rabbits.

We first explore the allometries for each block. The allometric change was more pronounced among basicranial bones, so indicating a comparatively “earlier development deacceleration” of facial bones, whose allometric growth has decreased, showing a modification of the typical times at which its developmental change takes place. These allometric differences can be attributed to retardation of face development, whilst the rest of the skull, i.e. the basicranial skeleton continues to grow. These features, due to retardation, implied significative asymmetry in sides for both splanchnocranial and basicranial bones, especially on the former, as an early cessation of development would disrupt the perfect right-left modelling. Asymmetries were located mainly on holes, passages for nerves and vessels and so of special functional importance. These results illustrate how artificially driven selection can lead to functional changes that decrease functional efficiency.

Moreover, our study also demonstrates a significant relationship between blocks when size is not considered, e.g., a correlated variation, resulting in shape proportional changes between them. Although cranial base usually reaches adult size before the face, in paedomorphic animals the face stops earlier its growth, breaking the statement that and the craniofacial elongation is the common aspect of postnatal growth in mammals (Cardini and Polly, 2013). If the face, that usually differentiates far later than the neurocranium, stops its normal development earlier in paedomporphic animals, this integration can produce more pronounced phenotypic differences between splanchnocranium and basicranium, as, ultimately, it imposes a constraint on cranial heterochrony. There is conserved the correlation between basicranium and splanchnocranium, but this has been reduced.

This study brings important new elements of major interest for discussing the role of the relationship between facial and basicranial shapes in the setting up of current extreme selective traits. It is one the first steps in a series of investigations on skull morphological variations in extreme paedomorphic selection. Future work will expand on the present analyses, and include other parts of the cranium and mandible to clarify the role and the importance of the size and shape of neotenic domestic mammals.

## Supporting information

The contents of all supporting data are the sole responsibility of the author.

## Conflicts of interest

The author declares no conflicts of interest to disclose related to this research.

## Acknowledgements

Author wishes to thank all the facilities given by CUNIPIC, who provided all corpses.

## Notes

### Competing Interest Statement

The authors have declared no competing interest.

